# Peramivir, an anti-influenza virus drug, exhibits potential anti-cytokine storm effects

**DOI:** 10.1101/2020.07.13.201806

**Authors:** Chen-xi Zhang, Ye Tu, De-pei Kong, Yinghua Li, Da-gui Chen, Wan-nian Zhang, Li Su, Chun-lin Zhuang, Zhi-bin Wang

**Affiliations:** Institute of Translational Medicine, Shanghai University, Shanghai, China; Department of Medicine, Shanghai East Hospital, Tongji University, 200120 Shanghai, China; School of Pharmacy, Second Military Medical University, 200433 Shanghai, China; School of Pharmacy, Ningxia Medical University, 750004 Yinchuan, China

**Keywords:** cytokine storm syndrome, COVID-19, peramivir, acute lung injury, multi-cytokines

## Abstract

Coronavirus Disease 2019 (COVID-19) infected by Severe Acute Respiratory Syndrome Coronavirus −2 (SARS-CoV-2) has been declared a public health emergency of international concerns. Cytokine storm syndrome (CSS) is a critical clinical symptom of severe COVID-19 patients, and the macrophage is recognized as the direct host cell of SARS-CoV-2 and potential drivers of CSS. In the present study, peramivir was identified to reduce TNF-α by partly intervention of NF-κB activity in LPS-induced macrophage model. In vivo, peramivir reduced the multi-cytokines in serum and bronchoalveolar lavage fluid (BALF), alleviated the acute lung injury and prolonged the survival time in mice. In human peripheral blood mononuclear cells (hPBMCs), peramivir could also inhibit the release of TNF-α. Collectively, we proposed that peramivir might be a candidate for the treatment of COVID-19 and other infections related CSS.

**Graphic Abstract:** 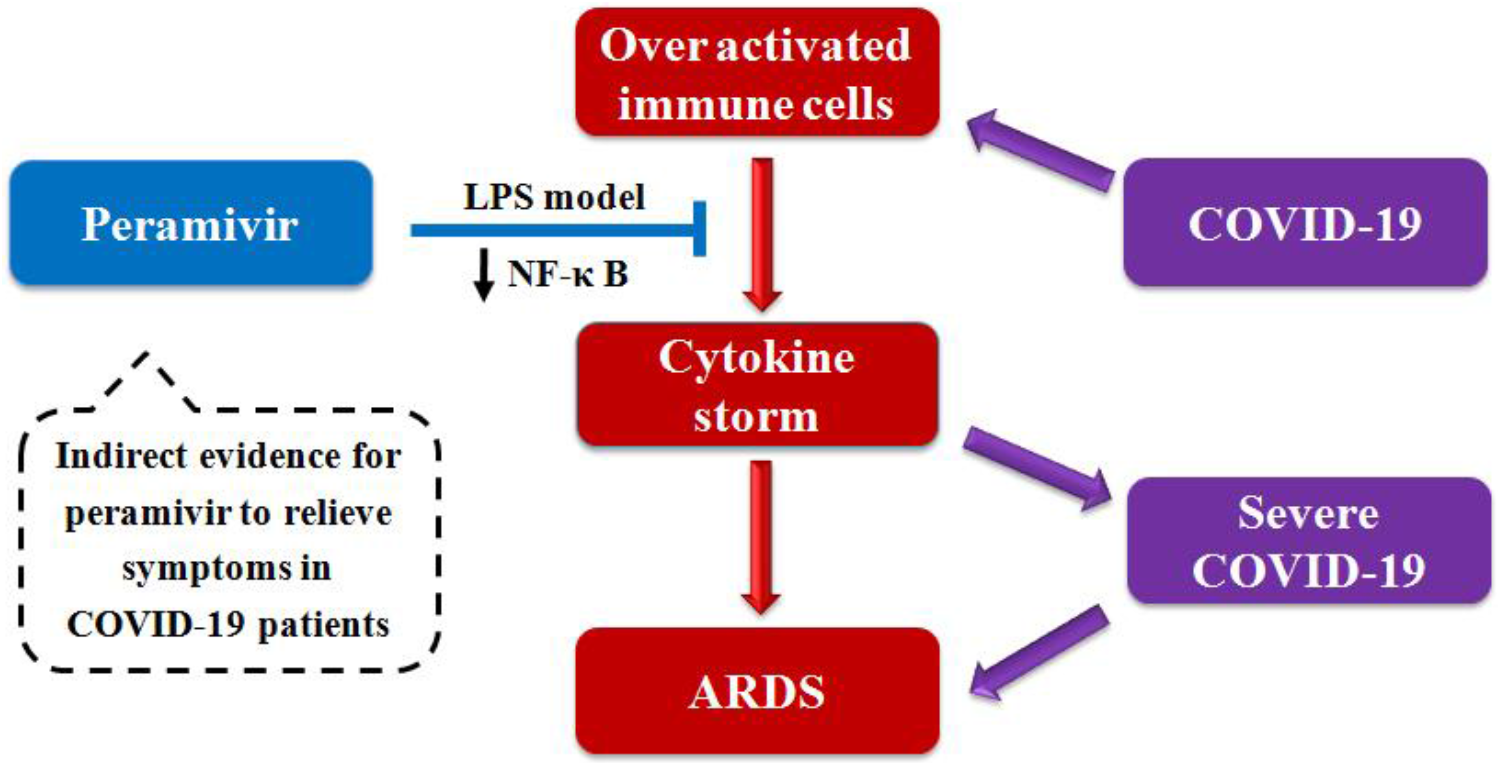

## INTRODUCTION

Coronavirus disease 2019 (COVID-19) caused by Severe Acute Respiratory Syndrome Coronavirus −2 (SARS-CoV-2) has been reported to infect more than 12 million people worldwide killing 500 thousand people (updated on Jul 10^th^, 2020).^1^ World Health Organization (WHO) has declared COVID-19 as a public health emergency of international concerns.^2^ COVID-19 patients show typical clinical symptoms of fever, fatigue, dry cough and pneumonia.^3–5^ Excessive inflammatory response leads to acute respiratory distress syndrome (ARDS), coagulopathy, and septic shock which can be fatal in critical cases.^3^ Gross anatomy identifies the main pathological features including exudation and hemorrhage, epithelium injuries, infiltration of inflammatory immune cells and fibrosis in the lungs of fatal patients.^3, 6–10^

A syndrome with a distinct cytokine storm was showed in a subgroup of patients with severe COVID-19, which has also been reported in SARS-CoV infected patients.^11–14^ The cytokine storm refers to an uncontrolled excessive inflammatory response, spreading from a local lesion to the whole body through the systemic circulation.^15, 16^ The plasma concentrations of inflammation related cytokines such as interleukins (IL) −2, −6, −7, and −10, tumor necrosis factor-α (TNF-α), interferon-γ-inducible protein 10 (IP10), granulocyte-colony stimulating factor (G-CSF), monocyte chemoattractant protein −1 (MCP-1), and macrophage inflammatory protein 1 alpha (MIP-1α) were significantly increased in the COVID-19 patients.^3, 17, 18^ Specially, the activation of alveolar macrophages was a characteristic abnormality.^8^

The leading cause of mortality is thought to be ARDS-induced respiratory failure, and patients generally receive supportive management in clinic practice.^17^ However, there is still no well-accepted effective treatment for COVID-19. The development of novel therapeutics has been mainly focused on antivirals^19, 20^ and vaccines.^11^ In addition, the cytokine storm is increasingly being recognized as a key node for the patients deteriorating to severe COVID-19.^3, 11, 17^ Therefore, anti-inflammatory therapy has been considered as one of appropriate clinical adjuvant treatment options, and the treatments include steroids (e.g., prednisone),^21^ selective cytokine blockade (e.g., tocilizumab),^11^ JAK inhibition (e.g., Baricitinib),^22^ intravenous immunoglobulin, Chinese medicines and blood purification.^11, 21^

Early in Feb, 2020, 75 of 99 COVID-19 patients received antiviral treatment including oseltamivir.^23^ Besides, oseltamivir is noted to have been widely used for confirmed or suspected COVID-19 cases in hospitals in China^24^ and Thailand (NCT04303299). The other two clinical trials (NCT04261270 and NCT04255017) involving oseltamivir in the treatment of COVID-19 are currently ongoing. However, the FDA-approved neuraminidase inhibitors including oseltamivir, zanamivir and peramivir were ineffective against the SARS-CoV-2 virus in vitro.^25^ There has been no exact evidence to date that oseltamivir is effective in the treatment of COVID-19 in clinic.^19^

It is reported that oseltamivir exhibited the antiviral activity of reducing pulmonary viral load, thereby the cytokines production was suppressed.^26^ Macrophages play a vital role in both SARS-CoV-2 virus -induced lung lesions and the host cytokine-mediated response.^8^ In our previous study, entecavir, a hepatitis B virus (HBV) inhibitor, was demonstrated to directly inhibit the release of cytokines in lipopolysaccharide (LPS) -stimulated macrophage model,^27^ which is a classic in vitro model to evaluate the anti-inflammatory activity of the drugs.^28^ Herein, we also examined whether these three neuraminidase inhibitors could inhibit the expression of inflammatory cytokines in LPS-stimulated macrophages in the present study. The results showed peramivir had the best ability to inhibit TNF-α by ~70% among the three compounds. Furthermore, we estimated the anti-cytokine storm effect and lung protection of peramivir in vivo. The anti-inflammatory effect of peramivir in human peripheral blood mononuclear cells (hPBMCs) was also observed.

## MATERIALS AND METHODS

### Materials

Compounds were purchased from TargetMol with a purity of > 98% (TargetMol). LPS (*E*. *coli* 0111:B4) was obtained from MilliporeSigma.

### Animal

Male C57BL/6J mice (18-22 g) were purchased from the Changzhou Cavens Laboratory Animal Co., China. All mice were kept under an automated 12 h dark-light cycle at a controlled temperature of 22°C ± 2°C and a relative humidity of 50%– 60% with free access to standard dry diet and tap water. All animal experiments were carried out in adherence with the NIH *Guide for the Care and Use of Laboratory Animals* (National Academies Press, 2011) and were approved by the Second Military Medical University Committee on Animal Care (EC11-055).

### CCS model

CCS was induced by a single i.p. injection of LPS (15 mg/kg), as described previously.^29^ After 4 h, mice were sacrificed and serum was collected. After 8 h, the left bronchus was ligated and 1 mL saline was perfused into right lung lobe to collect BALF, and the left lung was fixed with paraformaldehyde for histological analysis. Serum and BALF were further used for multi-cytokine analysis.

To investigate the effect of drugs on the survival time of CSS model mice, a single i.p. injection of saline (n=10), peramivir (20 and 60 mg/kg, n=10) was performed at 1 h before i.p. injection of a lethal dose of LPS (30 mg/kg) to mice, respectively. After modeling, mice survival was recorded every 2 h until 40 h.

### Preparation of the peritoneal macrophages

The peritoneal macrophages were obtained from the mice after i.p. injection of 3 ml of 3% thioglycolate as described previously.^29^ Briefly, the mice were sacrificed, and the macrophages were isolated by lavage with 5 ml of RPMI 1640 (Gibco Life Technologies), washed twice with PBS after 4 h of adherence, cultured in RPMI at 37°C and 5% CO_2_, and finally stimulated with 100 ng/ml LPS to harvest supernatants. The isolated cells were used for cytokine analysis and cell viability assays.

### Preparation of the hPBMCs

hPBMCs were obtained from freshly collected buffy coat fractions from healthy donors at the Tongren Hospital Affiliated to Shanghai Jiaotong University, China. Briefly, hPBMCs were isolated by centrifugation over a Ficoll-Paque (Pharmacia, Uppsala, Sweden) density gradient at 800 g for 20 min at room temperature in a Sorvall RT6000B (DuPont, Wilmington, DE, USA). Most hPBMC isolates were adherence cells that mainly contained macrophages and monocytes. Isolated hPBMCs were cultured in RPMI 1640, 100 U/mL penicillin–streptomycin (Invitrogen Life Technologies), and 10% heat-inactivated fetal calf serum (Gibco Life Technologies). 3 × 10^5^ cells were seeded in 96-well plates and incubated for 24 h at 37°C in a humidified atmosphere containing 5% CO_2_. hPBMCs were pretreated with peramivir at the concentrations of 2.5, 5 and 10 μM at 1 h before LPS (100 ng/ml) stimulation, and the supernatants were harvested at 6 or 12 h after LPS stimulation for cytokine analysis.

### Cell viability assay

Cell Counting Kit-8 (TY0312, Dojindo Molecular Technology, Japan) was used to measure cell viability. Briefly, 10 μL of CCK-8 solution was added, and cells were incubated for 1 h at 37°C. Absorbance was measured using a Cytation 5 Cell Imaging Multi-mode Reader (BioTek Instruments, USA) at a wavelength of 450 nm.

### Anti-inflammatory activity screening

We chose an in vitro model of LPS-stimulated peritoneal macrophages to induce TNF-α secretion, and screened potential anti-inflammatory molecules in the antiviral and antibacterial drug library. Briefly, 100 ng/ml LPS stimulated peritoneal macrophages for 4 h with simultaneous incubation of compounds at a concentration of 10 μM. The cell supernatant was diluted 10-fold and the TNF-α content was measured with a mouse TNF-α Elisa kit obtained from Invitrogen. The remaining cells were subjected to CCK8 assay to detect cytotoxicity.

### NF-κB luciferase activity assay

RAW264.7 cells stably transfected with an NF-κB-responsive luciferase construct, kindly provided by Prof. An Qin (Shanghai Jiaotong University, China), were seeded in 96-well plates at a density of 2 × 10^5^ cells per well, as previously described.^30^ After 24 h, cells were pretreated with peramivir for 1 h and stimulated with LPS for 6 h. Cells were dealt with a luciferase assay system (Promega) and the luciferase activity was calculated using a Cytation 5 Cell Imaging Multi-mode Reader (BioTek Instruments, USA).

### Western Blotting

Protein samples were separated by 10% SDS-PAGE, transferred to NC membrane and blocked with 5% non-fat milk in TBST. The membranes were washed with TBST and then incubated with the primary antibody for 6 hours at 4 degrees Celsius. The primary antibodies (1:1000) used were all from Cell Signaling Technology, USA and listed as follows: GAPDH antibody(#2118), stat3 antibody(#12640), phospho-stat3 antibody(#98543), SAPK/JNK antibody(#9252), phospho-SAPK/JNK antibody(#4668), p65 antibody(#4764), phospho-p65 antibody(#3033), IKKα antibody(#2682), phospho-IKKα/β antibody(#2697), IκBα antibody(#4812), phospho-IκBα antibody(#2859), p38 MAPK antibody(#8690), phospho-p38 MAPK antibody(#4511), Erk1/2 antibody(#4695) and phospho-Erk1/2 antibody(#4370). Then, the membranes were incubated in HRP-linked goat anti-rabbit IgG Antibody (1:10000, Cell Signaling Technology, USA, #7074) at room temperature for 1 hour and signals were detected by chemiluminescence (Bio rad, USA).

### ELISA

TNF-α released by mouse peritoneal macrophages was measured by a Mouse TNF-α ELISA Kit (Invitrogen, BMS607-3TEN) according to the manufacturer’s protocol. TNF-α released by hPBMCs were measured by Human TNF-α ELISA Kit (Youda, 1117202) according to the manufacturer’s protocol.

### Multi-cytokine measurement

The serum levels of a total of 12 virus-related cytokines were measured by a bead-based immunoassay panel (Mouse Anti-Virus Panel, Cat No: 740622, Biolegend, USA). The BALF levels of a total of 12 inflammatory cytokines were measured by a bead-based immunoassay panel (Mouse Inflammation Panel, Cat No: 740446, Biolegend, USA) on CytoFLEX Flow Cytometer (Beckman Coulter, USA) according to the manufacturer’s protocol.

### Immunofluorescence staining

Isolated peritoneal macrophages in eight-well LabTek slides (PEZGS0816, Millipore, Billerica, Massachusetts, USA) were pretreated with peramivir one hour before cell were simulated by LPS at a concentration of 1μg /ml for 30 min. Then cells were fixed in 4% paraformaldehyde, blocked with 0.4% Triton X-100/2% bovine serum albumin at room temperature for 1 h, and then incubated with primary antibodies for p65 (CST, #8242, 1:400 dilution) overnight at 4 °C. After being washed with PBST 3 times, the samples were incubated with Alexa Fluor 488 (Beyotime, A0423, 1:500 dilution) for 1 h and washed again with PBS. Nuclei were stained with DAPI. Images were obtained by confocal microscopy (TCS SP5, Leica, Solms, Germany).

### Histological analysis

The left lung lobes of mice were fixed using formalin, and then the tissues were paraffin-embedded Sections (5 μm) of formalin-fixed tissues were stained with haematoxylin and eosin (H&E) according to the manufacturer’s instructions, and were photographed with a microscope (Olympus Corporation, Tokyo, Japan). The histological characteristics of the lung injury (including alveolar edema and hemorrhage, the number of infiltrating leukocytes, and the thickness of the alveolar wall and epithelium) were evaluated. Each histological characteristic was evaluated on a scale of 0 to 3 (0, normal; 1, mild; 2, moderate; 3, severe).

### Statistical analysis

Data were expressed as means ± SEM. Statistical analyses used Student’s *t*-test, two-way ANOVA or Kaplan-Meier Survival Analysis. GraphPad software was used for data analysis. Statistical significance was indicated as follows: \**P* < 0.05, \*\**P* < 0.01, \*\*\**P* < 0.001, n.s. not significant.

## RESULTS

### Peramivir is an active anti-inflammatory agent without apparent cytotoxicity

The three neuraminidase inhibitors (peramivir, oseltamivir, and zanamivir, Fig. 1a) were explored for their ability to inhibit TNF-α-induced by LPS in macrophages. They inhibit the elevation of TNF-α by 67.2%, 35.6% and 13.1% at 10 μM, respectively (Fig. 1b). Furthermore, peramivir dose-dependently inhibited TNF-α release with the half-maximal inhibitory concentration (IC_50_) as 4.3 μM (Fig. 1c). Given that the inhibitory effect of the drugs on TNF-α might be achieved by cytotoxicity, we tested the cytotoxicity of peramivir in macrophages by a CCK-8 assay. It was demonstrated that no apparent toxicity was observed in the peramivir-treated macrophages at concentrations up to 40 μM (Fig. 1d).

**Fig. 1.**
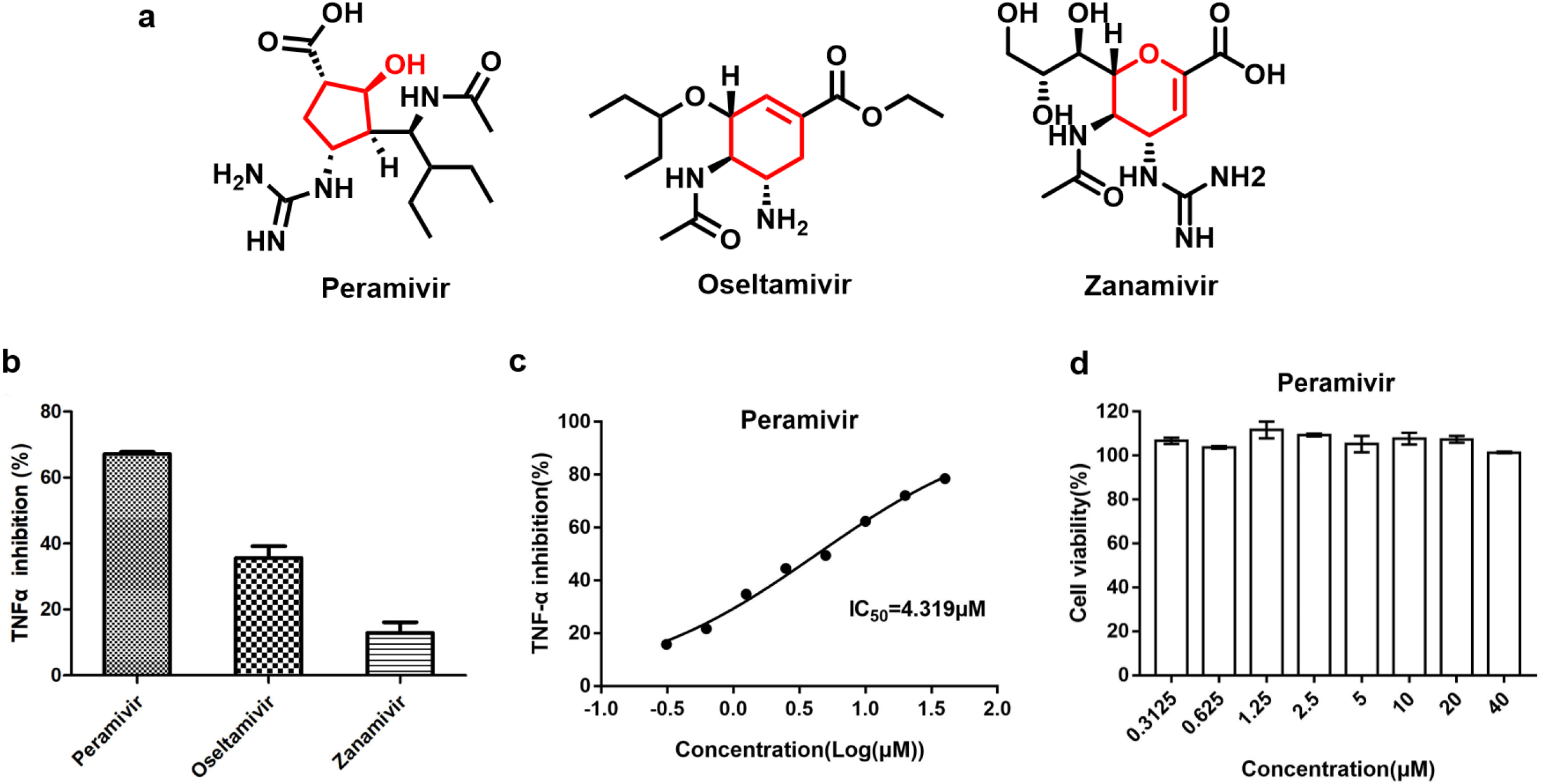
Identification of peramivir as anti-inflammatory agents. **a** Chemical structures of peramivir, oseltamivir and zanamivir. **b** Peramivir showed the strongest TNF-α inhibitory effect compared with oseltamivir and zanamivir. **c** The dose-response curves for the TNF-α inhibitions of peramivir exhibited IC_50_s of 4.3 μM. **d** Cell viabilities of macrophages with peramivir treatment at different concentrations.

### Peramivir inhibits LPS-induced cytokine storm in mice

We used LPS-induced cytokine storm syndrome (CSS) mouse model to evaluate the in vivo inflammatory inhibitory activity. Peramivir was administrated intraperitoneally (i.p.) at the concentration of 60 mg/kg at 1 h before LPS injection, and serum was collected at 4 h after LPS injection for further experiment. Twelve cytokines in total were simultaneously measured using a mouse antivirus panel by flow cytometric bead array. Compared with the control group, 7 cytokines including TNF-α, IL family (IL-6, IL-1β, IL-12), chemokines (MCP-1), interferon family (IFN-α, γ) were significantly decreased by the treatment (Table 1 and Fig. S1). The other 5 cytokines including IL-10, IP-10, chemokine (C-C motif) ligand 5 (CCL-5), granulocyte-macrophage colony stimulating factor (GM-CSF), CXC chemokine ligand 1 (CXCL1) were slightly downregulated without statistical significance.

**Table 1.**
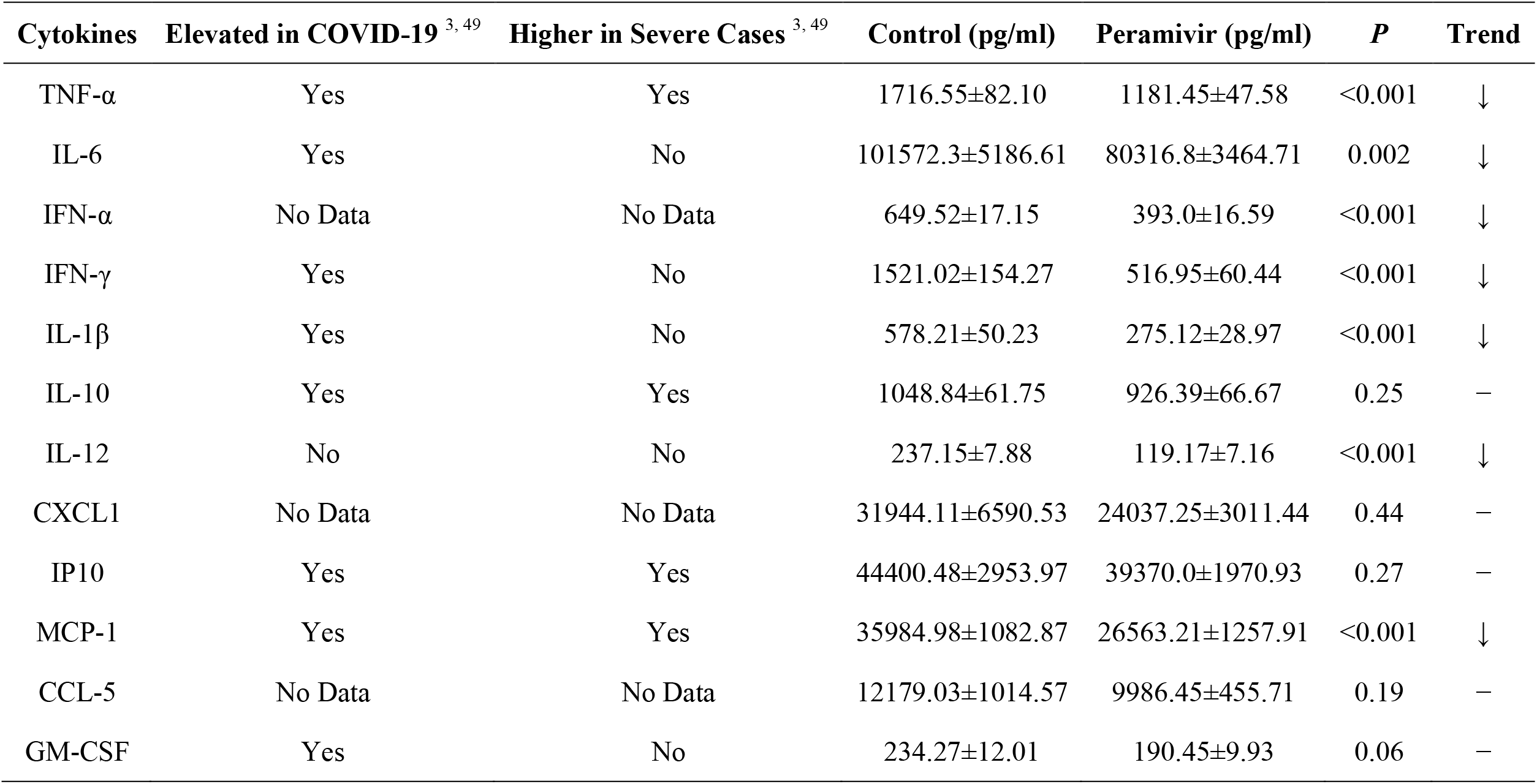
Clinical feature and experimental results of cytokines tested in mouse serum.

| Cytokines | Elevated in COVID-19<sup>3, 49</sup> | Higher in Severe Cases<sup>3, 49</sup> | Control (pg/ml) | Peramivir (pg/ml) | *P* | Trend |
| --- | --- | --- | --- | --- | --- | --- |
| TNF-α | Yes | Yes | 1716.55±82.10 | 1181.45±47.58 | <0.001 | ↓ |
| IL-6 | Yes | No | 101572.3±5186.61 | 80316.8±3464.71 | 0.002 | ↓ |
| IFN-α | No Data | No Data | 649.52±17.15 | 393.0±16.59 | <0.001 | ↓ |
| IFN-γ | Yes | No | 1521.02±154.27 | 516.95±60.44 | <0.001 | ↓ |
| IL-1β | Yes | No | 578.21±50.23 | 275.12±28.97 | <0.001 | ↓ |
| IL-10 | Yes | Yes | 1048.84±61.75 | 926.39±66.67 | 0.25 | — |
| IL-12 | No | No | 237.15±7.88 | 119.17±7.16 | <0.001 | ↓ |
| CXCL1 | No Data | No Data | 31944.11±6590.53 | 24037.25±3011.44 | 0.44 | — |
| IP10 | Yes | Yes | 44400.48±2953.97 | 39370.0±1970.93 | 0.27 | — |
| MCP-1 | Yes | Yes | 35984.98±1082.87 | 26563.21±1257.91 | <0.001 | ↓ |
| CCL-5 | No Data | No Data | 12179.03±1014.57 | 9986.45±455.71 | 0.19 | — |
| GM-CSF | Yes | No | 234.27±12.01 | 190.45±9.93 | 0.06 | — |

Given that cytokines in bronchoalveolar lavage fluid (BALF) could directly represent the inflammation status in the lungs,^7, 31^ we examined cytokines in BALF after 8 h of LPS stimulation. Twelve cytokines in total were simultaneously measured using a mouse inflammation panel by flow cytometric bead array. Compared with the control group, TNF-α and IL-6 were significantly decreased by the treatment (Table 2 and Fig. S2). IL-1β, IL-10, IL-17A and MCP-1 showed downregulation without significant difference.

**Table 2.**
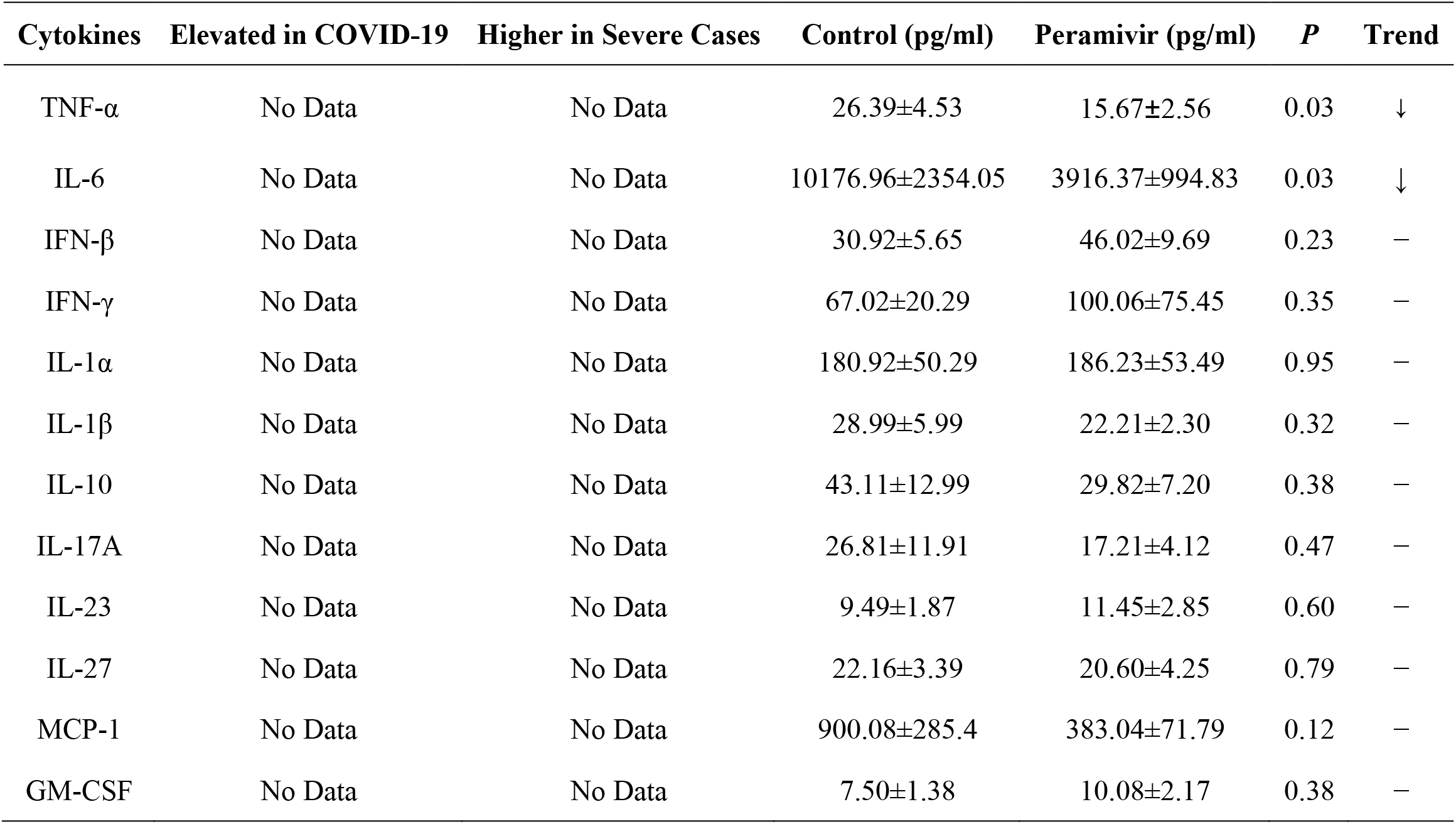
Clinical feature and experimental results of cytokines tested in mouse BALF.

| **Cytokines** | **Elevated in COVID-19** | **Higher in Severe Cases** | **Control (pg/ml)** | **Peramivir (pg/ml)** | ***P*** | **Trend** |
| --- | --- | --- | --- | --- | --- | --- |
| TNF-α | No Data | No Data | 26.39±4.53 | 15.67±2.56 | 0.03 | ↓ |
| IL-6 | No Data | No Data | 10176.96±2354.05 | 3916.37±994.83 | 0.03 | ↓ |
| IFN-β | No Data | No Data | 30.92±5.65 | 46.02±9.69 | 0.23 | — |
| IFN-γ | No Data | No Data | 67.02±20.29 | 100.06±75.45 | 0.35 | — |
| IL-1α | No Data | No Data | 180.92±50.29 | 186.23±53.49 | 0.95 | — |
| IL-1β | No Data | No Data | 28.99±5.99 | 22.21±2.30 | 0.32 | — |
| IL-10 | No Data | No Data | 43.11±12.99 | 29.82±7.20 | 0.38 | — |
| IL-17A | No Data | No Data | 26.81±11.91 | 17.21±4.12 | 0.47 | — |
| IL-23 | No Data | No Data | 9.49±1.87 | 11.45±2.85 | 0.60 | — |
| IL-27 | No Data | No Data | 22.16±3.39 | 20.60±4.25 | 0.79 | — |
| MCP-1 | No Data | No Data | 900.08±285.4 | 383.04±71.79 | 0.12 | — |
| GM-CSF | No Data | No Data | 7.50±1.38 | 10.08±2.17 | 0.38 | — |

### Peramivir effectively attenuates acute lung injury and prolong the survival in LPS-induced mice

The histological examinations of two COVID-19 death cases both showed alveolar damage with cellular fibromyxoid exudates, pulmonary edema and interstitial mononuclear inflammatory infiltrates.^7, 31^ The mice injected with LPS (i.p.) exhibited similar pathological features to ARDS, such as infiltration of inflammatory cells (black arrow), congestion (green arrow) and edema within thickened alveolar (yellow arrow) (Fig. 2a). In contrast, the alveolar structures of mice in the peramivir treated group were relatively intact, inflammatory cell infiltrations were significantly reduced, and mild alveolar thickening and less bleeding points or congestion were observed (Fig. 2a). Lung injury scores were calculated (Fig. 2b) to show significant protective effects of peramivir (score = 2.6 ± 0.6) to the lung tissues compared with that of control group (score = 4.8 ± 0.33). The survival time was prolonged in mice treated with peramivir in a dose-dependent manner after an i.p. injection of a lethal dose of LPS (30 mg/kg) compared with that in control mice (Fig. 2c).

**Fig. 2.**
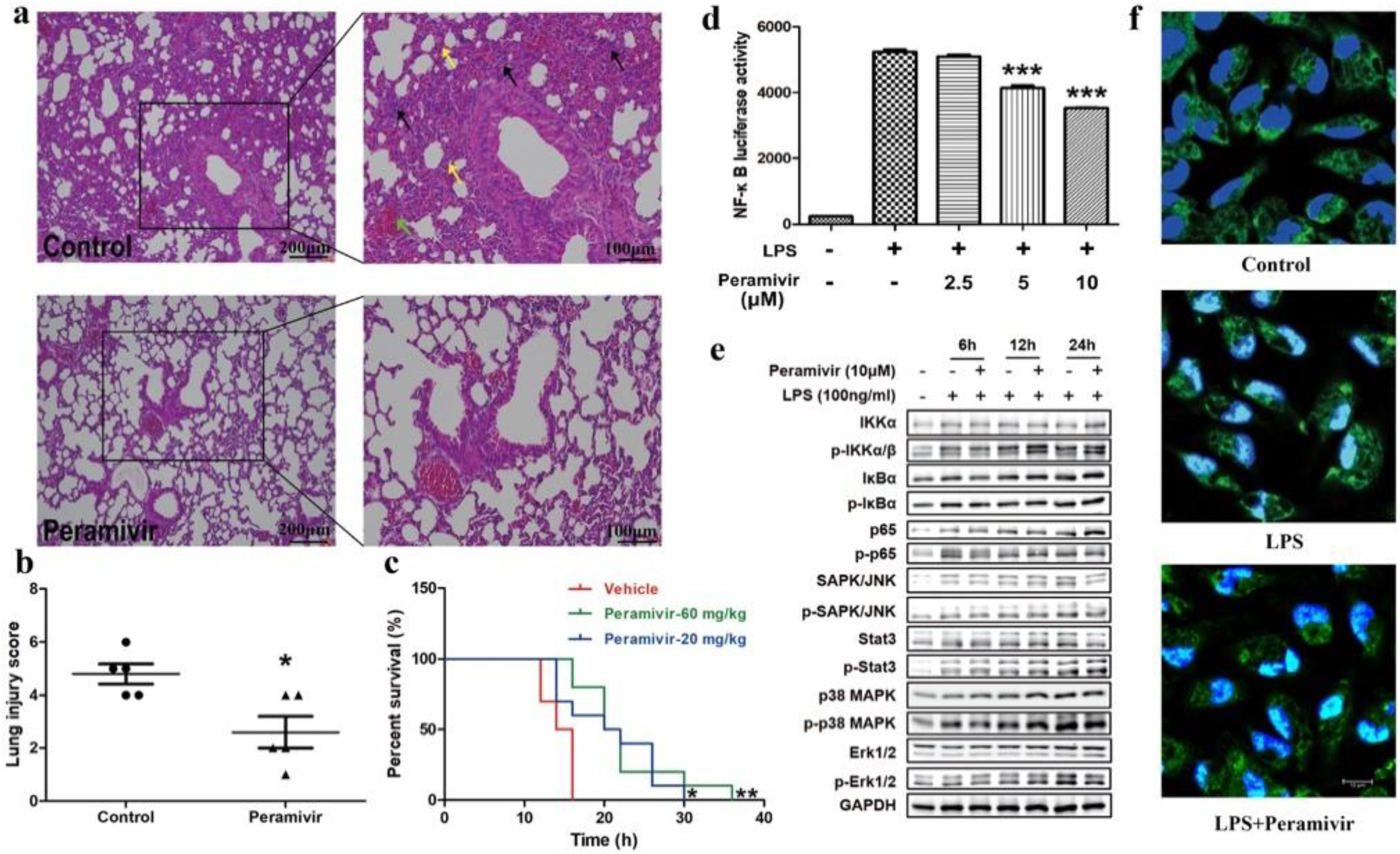
Peramivir effectively attenuates acute lung injury and prolong the survival in LPS-induced mice. **a** Representative images of lung H&E staining of control, and peramivir treatment groups. Black, green and yellow arrows indicated infiltration of inflammatory cells, congestion and edema within thickened alveolar, respectively. Scale bars, 100 or 200 μm as indicated. **b** Lung injury scores of control and peramivir treatment groups (n=5). \**P* < 0.05. **c** Survival time of LPS-induced CSS in control, and peramivir (20, 60 mg/kg) groups (n=10). Kaplan–Meier analysis was performed. \**P* < 0.05, \*\**P* < 0.01. **d** RAW264.7 cells were co-cultured with peramivir at concentrations of 2.5, 5 and 10 μM at 1 h before LPS stimulation. The activity of NF-κB luciferase was upregulated in all groups after 8 h, and there was a significant decline in cells co-cultured with peramivir in a dose-dependent manner. \*\*\**P* < 0.001. **e** The activation of the NF-κB, MAPK and STAT pathway in LPS-stimulated macrophages after the treatment of peramivir. **f** p65 nuclear translocation in LPS-stimulated macrophages after the treatment of peramivir (blue, DAPI; green, p65; cyan, cyan). Scale bars, 10 μm as indicated.

### Peramivir decreases NF-κB transcriptional activity in RAW264.7 and the peritoneal macrophages

NF-κB is an important transcriptional regulator in cells that participated in inflammatory responses, of which the activation induces the expression of multiple genes and production of cytokines consequently leading to cytokine storm.^32^ Peramivir dose-dependently inhibited LPS-induced NF-κB transcriptional activity in the RAW264.7 cells with a NF-κB reporter luciferase system (Fig. 2d). Furthermore, we detected some key factors in inflammation responses by western blot and found that the LPS-induced activation of NF-κB pathway (phosphorylation of p65) and MAPKs (phosphorylation of p38 and Erk1/2) were inhibited after peramivir intervention (Fig. 2e). Immunofluorescence images of LPS-stimulated peritoneal macrophages treated with or without peramivir indicated that LPS-induced nuclear translocation of p65 was attenuated by peramivir (Fig. 2f).

### Peramivir inhibits multi-cytokine releases in LPS-induced human peripheral blood mononuclear cells (hPBMCs)

Considering the translational value of peramivir in clinical practices, the release of TNF-α was tested in LPS-induced hPBMCs, which were obtained from two healthy donors. Peramivir (Fig. 3a, b and Fig. S3) significantly counteracted the level of TNF-α at 6 h and 12 h in a time- and dose-dependent manner without apparent toxicity.

**Fig. 3.**
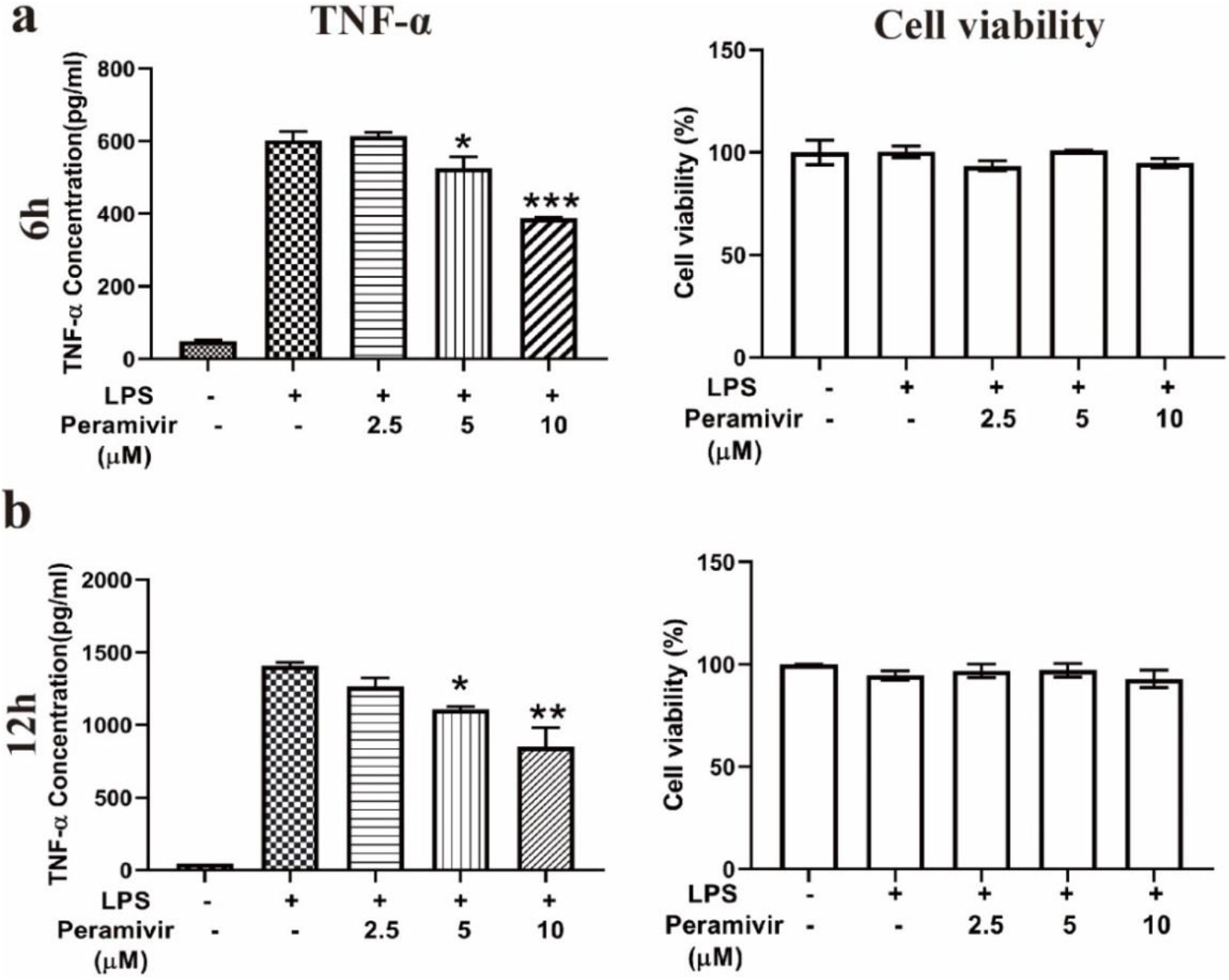
Peramivir inhibits cytokine release in LPS-induced hPBMCs from a health donor. TNF-α concentration was elevated by LPS stimulation. **a** and **b** Peramivir reduced TNF-α and IL-10 release in a time (6 and 12 h)- and dose (2.5, 5, 10 μM)-dependent manner. Peramivir showed no toxicity toward hPBMCs. \**P* < 0.05, \*\**P* < 0.01, *** *P* < 0.001.

## DISCUSSION

The three neuraminidase inhibitors bear similar pharmacophoric side chains (black, Fig. 1a) and different core scaffolds (red, Fig. 1a). Peramivir has a five-membered cyclopentanol ring, while oseltamivir and zanamivir have six-membered ring, which might be a critical part for the anti-inflammatory activity in the chemical structure of view.

Peramivir, an intravenous neuraminidase inhibitor, was approved for the emergency use in severe influenza in 2009 by the FDA. The antiviral effect of peramivir on influenza has been described previously,^33^ nevertheless, few studies have paid attentions to the anti-inflammatory activity of peramivir. The activation of the immune system is attributed to the virus-induced cytokine response.^34, 35^ Inhibition of these cytokines can potentially control the severity of the virus-induced inflammatory complications and finally reduce the mortality.^34, 35^ The inflammatory cytokines and influenza pathogenicity has been well correlated.^36, 37^ In mouse model of H1N1 influenza, peramivir inhibits the levels of TNF-α, IL6 and IFN-γ in the lung tissue.^38^ And compared to oseltamivir, peramivir shows more obvious anti-inflammatory effect.^39^ The anti-inflammatory effect of peramivir in vivo may be due to its antiviral symptomatic treatment, while we tried to investigate whether peramivir can directly inhibit the release of cytokines by inflammatory immune cells.

As the neuraminidase is not expressed in the SARS-CoV-2 virus, oseltamivir, zanamivir and peramivir were ineffective against the virus in vitro.^25^ In a clinical study, 75% of the COVID-19 patients received antiviral treatment including oseltamivir and 5 of them simultaneously infected with SARS-CoV-2 and influenza were recovered after treating with oseltamivir.^23^ We hypothesized that these neuraminidase inhibitors might have other effects including anti-inflammation, indicating the adjuvant therapeutic value of neuraminidase inhibitors in COIVD-19.

Viruses and bacteria induce immune cell activation and release of cytokines are Toll-like receptors (TLRs) dependent.^40^ The induction of inflammatory cytokines depends on the activation of NF-κB although they recognize different subtypes of TLRs.^41^ We assumed that SARS-CoV-2 may activate NF-κB on cytokine storm similar to that of SARS-Cov.^4^^1^ LPS activates immune cells such as monocytes and macrophages, causing the synthesis and release of inflammatory cytokines.^28^ TNF-α is one of the central cytokines involved in inflammation initiation and amplification in virus infections,^42^ and is reported to be elevated in critical COVID-19 cases,^3^ suggesting it as a proper indicator for in vitro drug screening.

Compared with oseltamivir, peramivir shows better inhibition of TNF-α in vitro. The reduction in TNF-α, IL-1β, IL-6, IL-12, IFN-α, IFN-γ and MCP-1 levels promoted by peramivir might play an important role in reducing stress due to the overactivated immune system and preventing organ damage of infected mice. This hypothesis was confirmed by pathological examination and lung index evaluation, which found that treatment with peramivir alleviated the severity of LPS-associated pneumonia and prolonged the survival time for mice. The lethal lung pathology caused by LPS was due to the excessive cytokine response that was primarily produced by the activated macrophages.^28^ Peramivir inhibited the levels of TNF-α and IL-6 in the BALF of LPS-induced mice. Furthermore, peramivir attenuated TNF-α induced by LPS both in mouse peritoneal macrophages and hPBMCs, which confirmed that peramivir can inhibit the inflammatory cytokine response mediated by macrophages.

In conclusion, the modulatory function of peramivir on LPS-induced inflammatory cytokines might contribute to the additional beneficial effect of the drug in antiviral therapy. This study provides evidence for the therapeutic value of peramivir for the potential application as an anti-inflammatory agent against cytokine dysregulation.

## ACKNOWLEDGEMENTS

We thank Profs Weiheng Xu and Pei Wang from Second Military Medical University, Drs Min Liu and Weiyuan Li from Tongren Hospital Affiliated to Shanghai Jiaotong University, China for the essential assistances with this study. We thank Taozhixing biotechnology (Shanghai, China) for providing us with the reagents and consumables urgently needed for the experiment. This work was supported in part by grants from the National Natural Science Foundation of China (81872880, 81703506 and 81703526) and the Young Elite Scientists Sponsorship Program by the China Association for Science and Technology (2017QNRC061).

## AUTHOR CONTRIBUTIONS

L. S., C.-L. Z., W.-N. Z, Z.-B. W. conceived and designed the experiments; L. S., Y. T., D.-G. C., C.-X. Z., Z.-B. W. participated in the experiments; L. S., Y. T., Z.-B. W. analyzed the data; L. S., C.-L. Z., Z.-B. W. wrote the manuscript; all the authors provided the final approval of the manuscript.

## Supplementary Figures

**Fig. S1.**
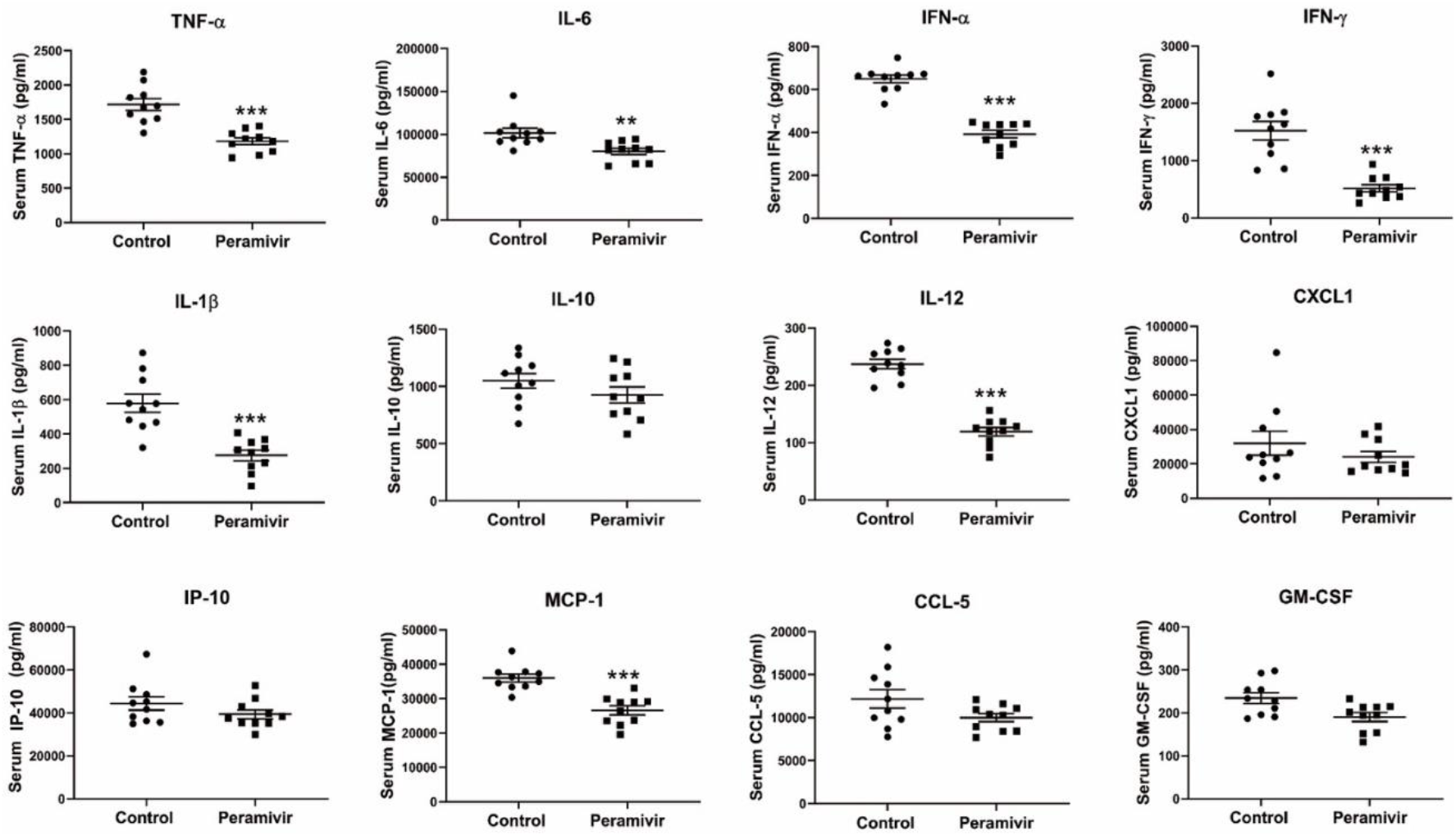
Effects of peramivir on different cytokines in mice serum. Compared with the control group, TNF-α, IL family (IL-6, IL-1β, IL-12), chemokines (MCP-1), interferon family (IFN-α, γ) were significantly decreased by the treatment. **P <0.01, ***P < 0.005.

**Fig. S2.**
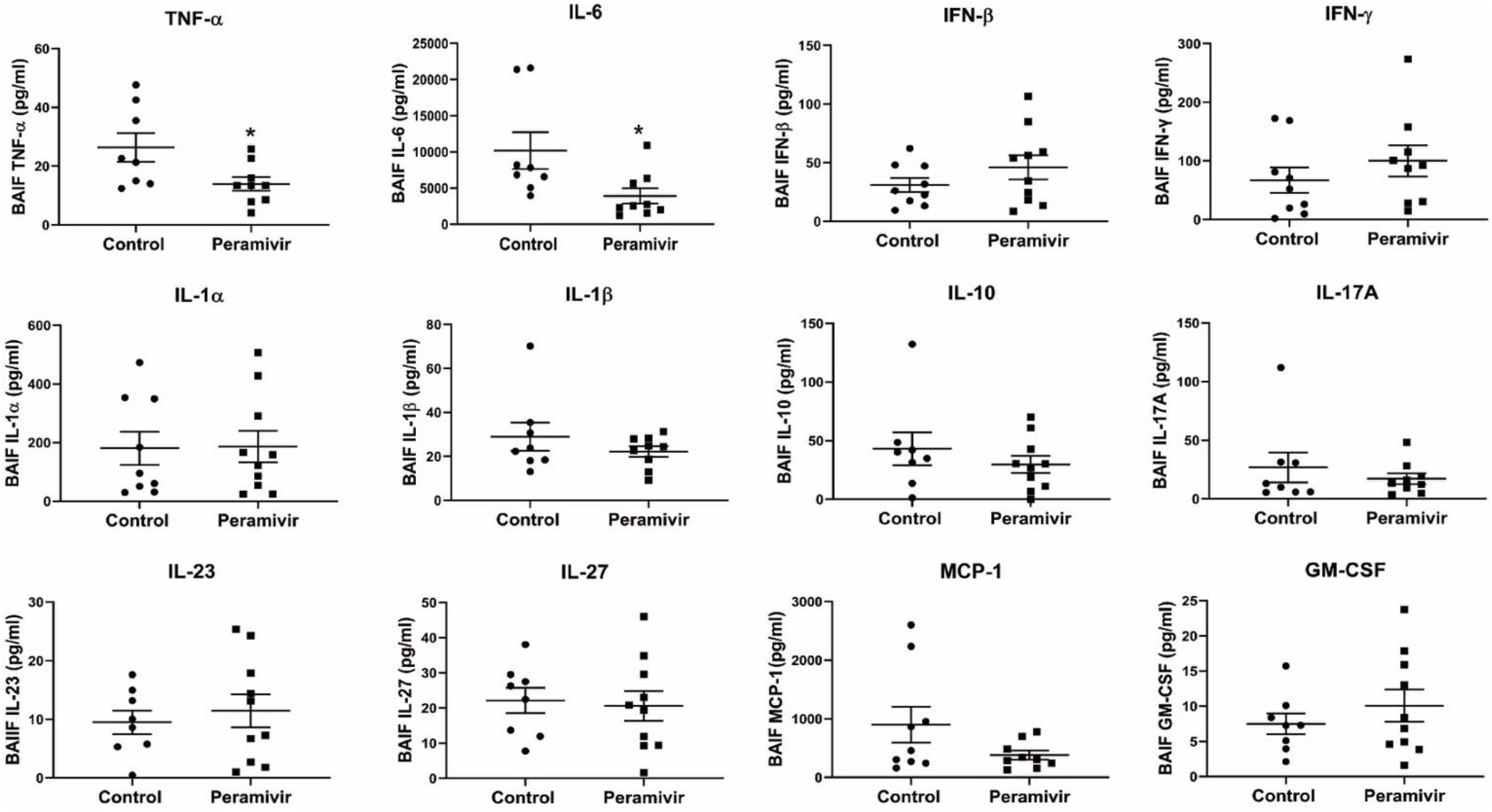
Effects of peramivir on different cytokines in mouse bronchoalveolar lavage fluid (BALF). Compared with the control group, TNF-α and IL-6 were significantly decreased by the treatment. *P < 0.05.

**Fig. S3.**
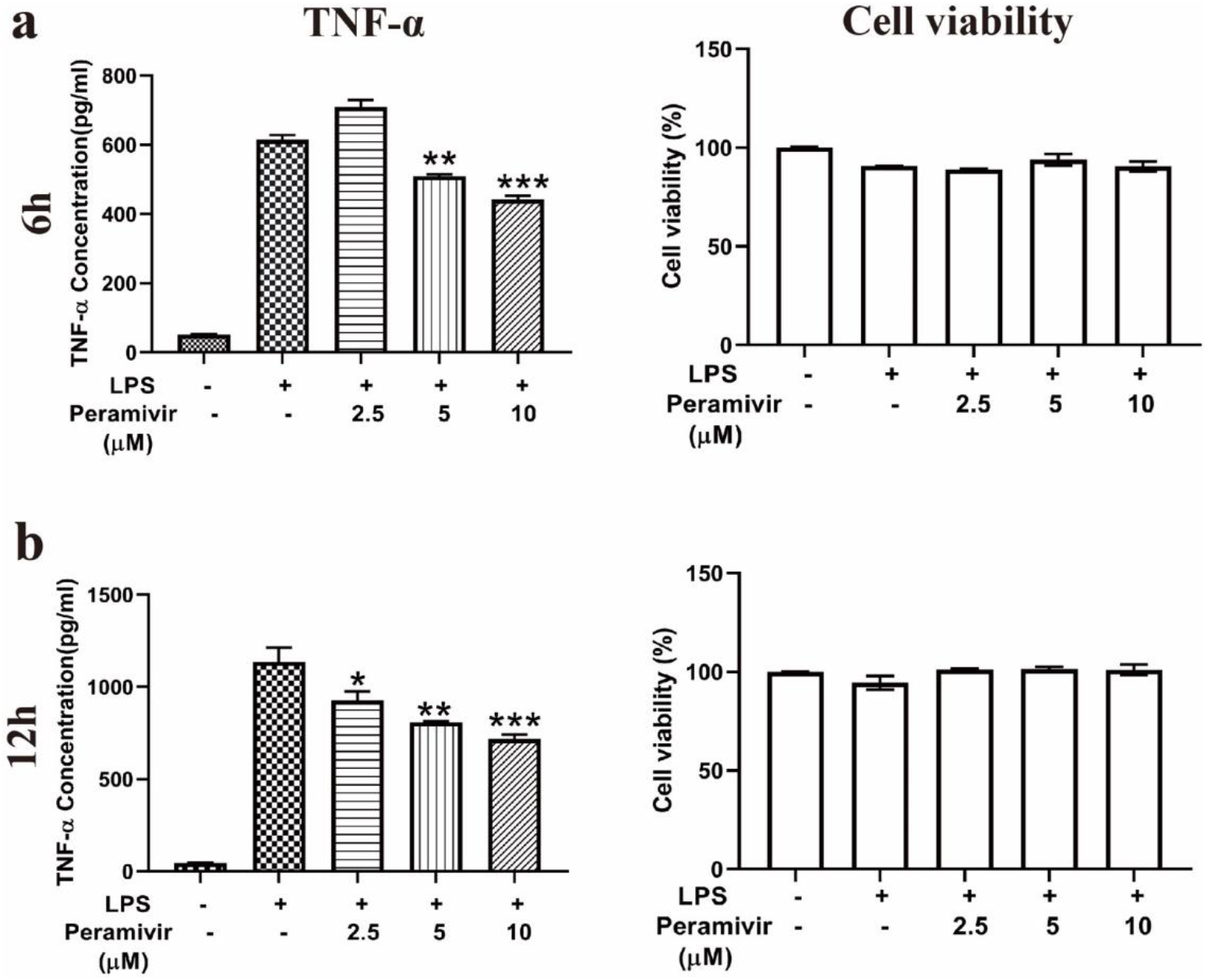
Peramivir inhibits TNF-α release in LPS-induced hPBMCs from the second health donor. TNF-α concentration was elevated by LPS stimulation. a and b Peramivir reduced TNF-α release in dose (2.5, 5, 10 μM)-dependent manner. Peramivir showed no toxicity toward hPBMCs. *P < 0.05, **P <0.01, ***P < 0.005.

